# Phylogenetically conserved peritoneal fibrosis response to an immunologic adjuvant in ray-finned fishes

**DOI:** 10.1101/2020.07.08.191601

**Authors:** Milan Vrtílek, Daniel I. Bolnick

**Author notes:** **Corresponding author:** The Czech Academy of Sciences, Institute of Vertebrate Biology, Květná 8, 603 65 Brno, Czech Republic.

## Abstract

Antagonistic interactions between hosts and parasites may drive the evolution of novel host defenses, or new parasite strategies. Host immunity is therefore one of the fastest evolving traits. But where do the novel immune traits come from? Here, we test for phylogenetic conservation in a rapidly evolving immune trait – peritoneal fibrosis. Peritoneal fibrosis is a costly defense against novel specialist tapeworm *Schistocephalus* solidus (Cestoda) expressed in some freshwater populations of threespine stickleback fish (*Gasterosteus aculeatus*, Perciformes). We asked whether stickleback fibrosis is a derived species-specific trait or an ancestral immune response that was widely distributed across ray-finned fish (Actinopterygii) only to be employed by threespine stickleback against the specialist parasite. We combined literature review on peritoneal fibrosis with a comparative experiment using either parasite-specific, or non-specific, immune challenge in deliberately selected species across fish tree of life. We show that ray-finned fish are broadly, but not universally, able to induce peritoneal fibrosis when challenged with a generic stimulus (Alum adjuvant). The experimental species were, however, largely indifferent to the tapeworm antigen homogenate. Peritoneal fibrosis, thus, appears to be a common and deeply conserved fish immune response that was co-opted by stickleback to adapt to a new selective challenge.

## INTRODUCTION

The comparative immunology research makes it clear that many of the fastest-evolving and most polymorphic genes in vertebrates are involved in immunity (Litman and Cooper 2007; Lazzaro and Clark 2012; Slodkowicz and Golman 2020). Most notable is the evolutionary reshuffling of the genes coding Toll-like receptors (TLR) (Solbakken et al. 2017; Velová et al. 2018), or the diversity of major histocompatibility complex (MHC) genes (Malmstrøm et al. 2016; Radwan et al. 2020). Conversely, the broad outlines of innate immunity are ancient, such as one of the most ancestral immune cytokines, transforming growth factor β (TGF-β), which seems to be conserved across the animal kingdom (Herpin et al. 2004). And yet, even some highly conserved immune genes and processes have been lost or changed past recognition in certain vertebrate clades, such as the loss of MHCII genes in the Atlantic cod (Malmstrøm et al. 2013). This contrast between deep evolutionary conservation, and rapid co-evolutionary dynamics, is puzzling. What features of the immune system are highly conserved, and what are evolutionarily labile?

Vertebrates possess, in principle, two functionally distinct strategies combining innate and adaptive immunity to cope with infection according to parasite type (Flajnik and Du Pasquier 2004; Allen and Maizels 2011). Type 1 immune response is triggered by fast reproducing pathogens, as microbes, with the aim to quickly eliminate the infection through pro-inflammatory trajectory (Allen and Maizels 2011). On the other hand, type 2 immune response is typically directed to reduce the effect of a multicellular parasite, such as a helminth worm, by containment and encapsulation (Allen and Maizels 2011; Gause et al. 2013). Type 2 immunity largely shares signaling pathways with tissue repair and wound healing (Gause et al. 2013; Thannickal et al. 2014). Perpetual tissue damage and wound healing may, however, result in excessive accumulation of fibrous connective matter called fibrosis (Thannickal et al. 2014). This fibrosis has been found to effectively suppress growth of certain parasites, or even lead to parasite death (Weber et al., in preparation). However, the benefits of tissue repair and parasite containment can come with a cost from chronic type 2 immune response during persistent or recurrent infections, which may develop into serious health issues or even death (Gause et al. 2013; De Lisle and Bolnick 2020). Here, we measure the extent of evolutionary conservation of a key immune phenotype, fibrosis.

Recent findings on inter-population variation in helminth resistance from threespine stickleback fish (*Gasterosteus aculeatus*) demonstrate that anti-helminthic fibrosis response is a fast-evolving immune trait (Weber et al. 2017a). Stickleback, originally a marine species, has only recently invaded freshwater habitats where it experienced greater risk of acquiring parasitic tapeworm *Schistocephalus solidus* (Cestoda) through feeding on freshwater copepods (Barber and Scharsack 2010; Rahn et al. 2016). When ingested, the tapeworm larva migrates through the intestinal wall to the peritoneal cavity of the fish and grows to its final size, often >30% the host’s mass (Arme and Owen 1967; Ritter et al. 2017). The threespine stickleback is the obligate intermediate host of this specialized parasite. Some populations of stickleback have evolved a capacity to suppress *S. solidus* growth by encapsulating it in fibrotic tissue, sometimes leading to successfully killing and eliminating the parasite (Weber et al. 2017b).

This presumably beneficial form of resistance has costs in greatly suppressing female gonad development and male reproduction (De Lisle and Bolnick 2020; Weber et al., in preparation). These costs may explain why the intensive peritoneal response to *S. solidus* has evolved in only some lake populations, and in some geographic regions of the sticklebacks’ range (Weber et al. 2017a). In the other populations, stickleback have apparently adopted a non-fibrotic tolerance response to reproduce despite infection (Weber et al. 2017a). These non-fibrotic populations exhibit active up-regulation of fibrosis-suppression genes in response to cestode infection (Lohman et al. 2017; Fuess et al. 2020). The ancestral marine populations come very rarely into contact with *S. solidus* which does not hatch in saline water (Barber and Scharsack 2010), and do not exhibit observable fibrosis in the wild or in captivity (Hund et al. 2020). These various marine and freshwater populations have been diverging only since Pleistocene deglaciation (~12,000 years), indicating that their fibrosis response has evolved surprisingly quickly for such a fundamental immune process.

The peritoneal fibrosis can reliably be provoked in both fibrotic and non-fibrotic populations of stickleback by a generalized immune challenge (injection with a non-specific Alum adjuvant) (Hund et al. 2020), while only the resistant populations initiate fibrosis in response to tapeworm protein injection. So, the physiological capacity to initiate fibrosis seems conserved in stickleback and its sensitivity to the tapeworm fast-evolving. We therefore wished to determine whether this peritoneal fibrosis is similarly labile, or conserved across a broader range of fish species. One possibility is that peritoneal fibrosis is unique to stickleback, which is the only fish species to host *S. solidus*, an unusual parasite in that it invades the fish peritoneal cavity (Barber and Scharsack 2010). Or, fibrosis may be a widely conserved trait that stickleback have uniquely co-opted to deal with this specialist parasite species. Is then peritoneal fibrosis an ancestral character state dating back to the origin of teleosts, or beyond?

To address this question, we begin with a broad literature review of fibrosis in fishes to systematically summarize documented instances for the first time. Then, we present an experimental test of how widely ray-finned fish (Actinopterygii) exhibit peritoneal fibrosis response towards an immune challenge based on a recent experimental study of fibrosis response in stickleback (Hund et al. 2020).

## METHODS

### Literature review

We performed search for articles mentioning fibrosis or encapsulation from Actinopterygii using Web of Science database. We then sorted the collected suitable articles according to the presence of fibrosis or encapsulation, its topology and cause. We also extracted the taxonomic information and present the results in a pivot table. For detailed methods, please see *Appendix*.

### Species selected for the experiment

To conduct a phylogenetically broad assay of peritoneal fibrosis response in ray-finned fishes (Actinopterygii), we deliberately chose a diverse set of species spread across the phylogenetic tree of Actinopterygii. We experimentally vaccinated 17 species of fish (Table 1). These species were chosen to achieve broad phylogenetic diversity, but were restricted to commercially available small fish (body length of 1-4”, or 2-10 cm, and live weight 0.2-6.0 g). We obtained most of the fish from a fish reseller. Local trout hatchery donated fingerlings of rainbow trout (*Oncorhynchus mykiss*). We received eggs of the turquoise killifish (*Nothobranchius furzeri*, population MZCS 222) from a stock retained at the Institute of Vertebrate Biology CAS in Brno, Czech Republic. We hatched and maintained the killifish according to breeding protocol (Polačik et al. 2016) until they reached two months when fully mature. We also included wild-caught threespine stickleback (*Gasterosteus aculeatus*) that came from two distinct populations (Loch Hosta and Loch a’ Bharpa), both originating from North Uist, Scotland, UK, provided by Andrew MacColl (University of Nottingham, UK) and maintained at our housing facility for six months before the experiment.

**Table 1.**
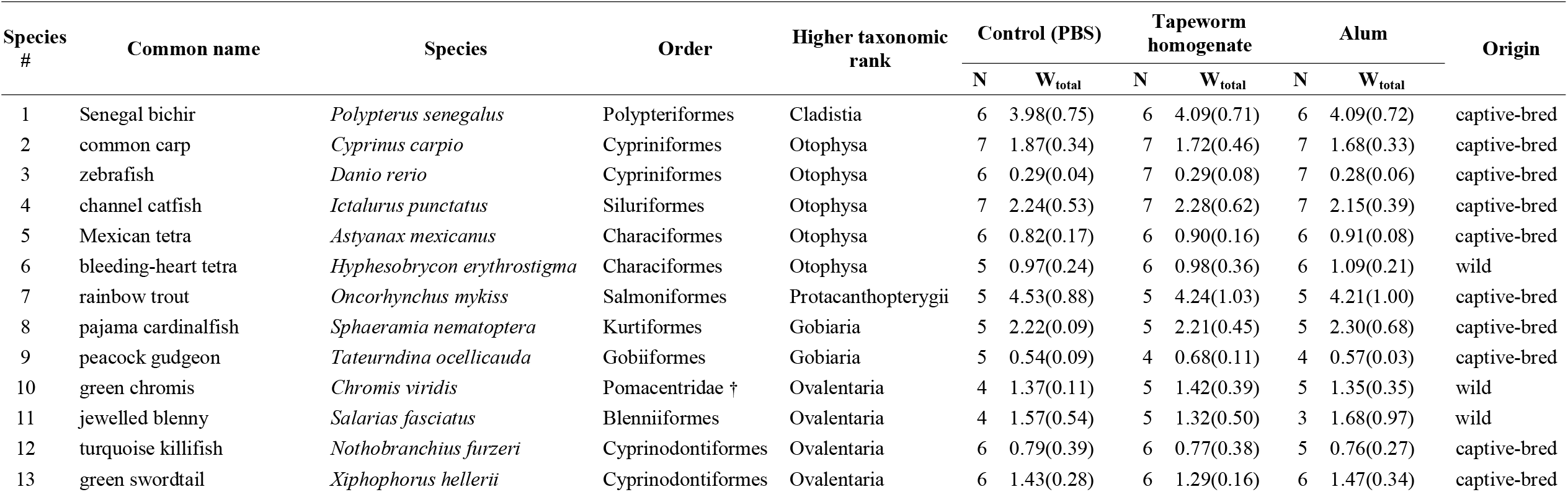
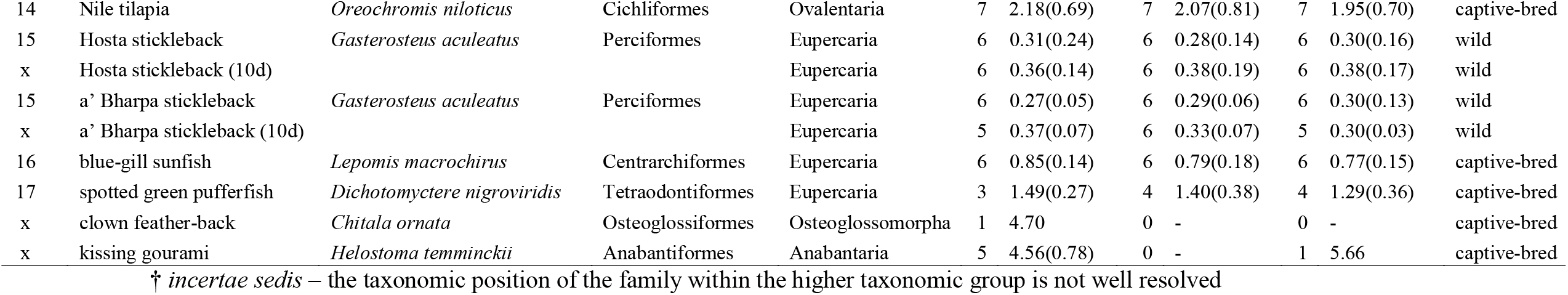
Species selected for the experimental test of peritoneal fibrosis response. We show the sample size, N, and average with standard deviation (SD) for live weight before dissection in grams, W_total_, in each experimental treatment. Note that Species # corresponds to species number in Fig. 2. We also present data for two species that we did not use in the analysis as well as for the stickleback that were kept in the treatment for 10 days (“10d”) indicated by “x”. The clown feather-back (*Chitala ornata*) were extremely fibrosed (level 3) before the experiment started. The kissing gourami (*Helostoma temminckii*) developed white spot disease during the experiment.

| species<br># | Common name | Species | Order | Higher taxonomic<br>rank | Control (PBS) |  | Tapeworm<br>homogenate |  | Alum |  | Origin |
| --- | --- | --- | --- | --- | --- | --- | --- | --- | --- | --- | --- |
| | | | | | N | $W_{\text{total}}$ | N | $W_{\text{total}}$ | N | $W_{\text{total}}$ | |
| 1 | Senegal bichir | *Polypterus senegalus* | Polypteriformes | Cladistia | 6 | 3.98(0.75) | 6 | 4.09(0.71) | 6 | 4.09(0.72) | captive-bred |
| 2 | common carp | *Cyprinus carpio* | Cypriniformes | Otophysa | 7 | 1.87(0.34) | 7 | 1.72(0.46) | 7 | 1.68(0.33) | captive-bred |
| 3 | zebrafish | *Danio rerio* | Cypriniformes | Otophysa | 6 | 0.29(0.04) | 7 | 0.29(0.08) | 7 | 0.28(0.06) | captive-bred |
| 4 | channel catfish | *Ictalurus punctatus* | Siluriformes | Otophysa | 7 | 2.24(0.53) | 7 | 2.28(0.62) | 7 | 2.15(0.39) | captive-bred |
| 5 | Mexican tetra | *Astyanax mexicanus* | Characiformes | Otophysa | 6 | 0.82(0.17) | 6 | 0.90(0.16) | 6 | 0.91(0.08) | captive-bred |
| 6 | bleeding-heart tetra | *Hyphesobrycon erythrostigma* | Characiformes | Otophysa | 5 | 0.97(0.24) | 6 | 0.98(0.36) | 6 | 1.09(0.21) | wild |
| 7 | rainbow trout | *Oncorhynchus mykiss* | Salmoniformes | Protacanthopterygii | 5 | 4.53(0.88) | 5 | 4.24(1.03) | 5 | 4.21(1.00) | captive-bred |
| 8 | pajama cardinalfish | *Sphaeramia nematoptera* | Kurtiformes | Gobiaria | 5 | 2.22(0.09) | 5 | 2.21(0.45) | 5 | 2.30(0.68) | captive-bred |
| 9 | peacock gudgeon | *Tateurndina ocellicauda* | Gobiiformes | Gobiaria | 5 | 0.54(0.09) | 4 | 0.68(0.11) | 4 | 0.57(0.03) | captive-bred |
| 10 | green chromis | *Chromis viridis* | Pomacentridae † | Ovalentaria | 4 | 1.37(0.11) | 5 | 1.42(0.39) | 5 | 1.35(0.35) | wild |
| 11 | jewelled blenny | *Salarias fasciatus* | Blenniiformes | Ovalentaria | 4 | 1.57(0.54) | 5 | 1.32(0.50) | 3 | 1.68(0.97) | wild |
| 12 | turquoise killifish | *Nothobranchius furzeri* | Cyprinodontiformes | Ovalentaria | 6 | 0.79(0.39) | 6 | 0.77(0.38) | 5 | 0.76(0.27) | captive-bred |
| 13 | green swordtail | *Xiphophorus hellerii* | Cyprinodontiformes | Ovalentaria | 6 | 1.43(0.28) | 6 | 1.29(0.16) | 6 | 1.47(0.34) | captive-bred |
| 14 | Nile tilapia | *Oreochromis niloticus* | Cichliformes | Ovalentaria | 7 | 2.18(0.69) | 7 | 2.07(0.81) | 7 | 1.95(0.70) | captive-bred |
| 15 | Hosta stickleback | *Gasterosteus aculeatus* | Perciformes | Eupercaria | 6 | 0.31(0.24) | 6 | 0.28(0.14) | 6 | 0.30(0.16) | wild |
| x | Hosta stickleback (10d) |  |  | Eupercaria | 6 | 0.36(0.14) | 6 | 0.38(0.19) | 6 | 0.38(0.17) | wild |
| 15 | a' Bharpa stickleback | *Gasterosteus aculeatus* | Perciformes | Eupercaria | 6 | 0.27(0.05) | 6 | 0.29(0.06) | 6 | 0.30(0.13) | wild |
| x | a' Bharpa stickleback (10d) |  |  | Eupercaria | 5 | 0.37(0.07) | 6 | 0.33(0.07) | 5 | 0.30(0.03) | wild |
| 16 | blue-gill sunfish | *Lepomis macrochirus* | Centrarchiformes | Eupercaria | 6 | 0.85(0.14) | 6 | 0.79(0.18) | 6 | 0.77(0.15) | captive-bred |
| 17 | spotted green pufferfish | *Dichotomyctere nigroviridis* | Tetraodontiformes | Eupercaria | 3 | 1.49(0.27) | 4 | 1.40(0.38) | 4 | 1.29(0.36) | captive-bred |
| x | clown feather-back | *Chitala ornata* | Osteoglossiformes | Osteoglossomorpha | 1 | 4.70 | 0 | - | 0 | - | captive-bred |
| x | kissing gourami | *Helostoma temminckii* | Anabantiformes | Anabantaria | 5 | 4.56(0.78) | 0 | - | 1 | 5.66 | captive-bred |

† *incertae sedis* – the taxonomic position of the family within the higher taxonomic group is not well resolved

### Fish housing

We standardized housing conditions across most of the species. We planned to have 5-7 individuals per species per treatment. We prepared 20-gal (~76 L) tanks with reverse-osmosis (RO) water conditioned to conductivity between 700-800 µS/cm with sea salt (Instant Ocean^®^). We adjusted salinity for marine species to 35 mg/kg. Each tank contained air driven sponge filter and heater with temperature set to 25 °C (water temperature ranged between 24.5-26 °C). We also provided a seaweed-like plastic shelter for fish. We fed fish every morning with frozen bloodworms (Chironomidae), mysis shrimp (Mysidae), or dried sushi nori seaweed (*Pyropia* sp.) according to species-specific diet requirements. We held the rainbow trout at water temperature of 12 °C as our standard temperature would be stressful to them. Similarly, stickleback are sensitive to high temperatures and we therefore kept them in their original recirculation system at 19 °C and 1900 µS/cm throughout the treatment. All the other fish were habituated to the common tank setup for at least five days before treatment.

### Reagents

Drawing on a recent experimental study of fibrosis response in stickleback (Hund et al. 2020), we challenged selected fish species with a generalized immune stimulant (Alum vaccination adjuvant), or with antigens from a specialist helminth parasite (*S. solidus*) that induces peritoneal fibrosis in some populations of threespine stickleback. Saline solution served as a control. All three treatments were delivered via peritoneal injection (the site of *S. solidus* infection), for all 17 species.

The control treatment was an injection of 1X phosphate-buffered saline (PBS) (20 µL per 1 g of species average of live weight), which was also the solution for delivering the other treatments. The second treatment consisted of tapeworm antigen homogenate (abbreviated TH) suspended in PBS. Hund et al. (2020) showed that injection of 9 mg of TH per 1kg live fish mass (0.009 mg/g) induced rapid fibrosis in tapeworm-resistant stickleback populations. We obtained *S. solidus* tapeworms dissected from wild-caught threespine stickleback (Gosling Lake, Vancouver Island, BC, Canada). We used two tapeworms to prepare the homogenate, sonified them in PBS on ice and then centrifuged the suspension at 4000 RPM at 4 °C for 20 minutes. We assessed overall protein concentration in the upper fraction of the solution using RED 660™ protein assay (G-Biosciences) measured in triplicates and then diluted the sample to 0.45 mg/mL. We aimed at injecting 20 µL of the solution per 1 g of live fish weight and to obtain the desired dose 0.009 mg of tapeworm protein for 1 g of fish weight (or 9 mg/kg, as in Hund et al. (2020)). We then aliquoted the homogenate in 0.6-ml Eppendorf tubes and stored at −20 °C for later injections. Using the tissue of *S. solidus*, we wanted to test whether the stickleback response to this specialized parasite is unique among other fishes. In principle, for instance, it is possible that *S. solidus* protein contains distinctive antigens that any fish species would recognize as foreign.

The third treatment was a 1% Alum solution (20 µL per 1 g of species average of fish live weight). Alum promotes activation of innate immune response and is commonly used as a vaccine adjuvant (Kool et al. 2012). We dissolved 2% AlumVax Phosphate (OZ Biosciences) in 1:1 with PBS. This concentration of Alum induces peritoneal fibrosis in marine and freshwater stickleback population irrespective of their tapeworm resistance (Hund et al. 2020).

### Injections

At their arrival to our fish facility, we weighed each fish species on a balance to 0.01 g (Gene Mate GP-600) to estimate total volume of solution to be injected per individual (based on species average live mass). We injected 20 µL of the solution per 1 g of average live weight with Ultra-Fine insulin syringes. We always filled syringes aseptically under laminar flow cabinet 1-2 days before the injections, stored them at 4 °C and used 1 syringe per individual. Prior work (Hund et al. 2020) confirmed that syringes prepared in this way were effective at inducing fibrosis. We injected fish peritoneally through their left flank after anesthesia with MS-222 (200 mg/L for up to two minutes). Fish were then allowed to recover from anesthesia in a highly aerated tank water. We returned them to their original tank once they were swimming upright, but typically after more than five minutes after the injection. The different treatment groups were held in separate tanks because peritoneal fibrosis is a specific response to the treatment unaffected by common housing conditions (Hund et al. 2020).

### Dissections and scoring fibrosis level

We euthanized fish five days post-injection, using an overdose of MS-222 (500 mg/L, > 5 min). Because trout and stickleback were kept at lower temperatures, which may slow immune responses (Rijkers et al. 1980), we dissected trout 10 days after injection, and stickleback at both 5 and 10 days. We also dissected 1-2 individuals from each species prior the injections to examine species-specific anatomy and assess the baseline level of peritoneal fibrosis before injections. We dissected fish immediately after euthanasia under stereo-microscope, photographed and scored their level of fibrosis, using an ordinal categorical score. The peritoneal fibrosis score ranged between 0 and 3. Zero represents the absence of noticeable fibrosis, where the internal organs (liver, intestine, gonads) move freely apart from each other and from the peritoneal wall when moved with tweezers. Level 1 was for organs adhered together forming an interconnected conglomerate that moves as a unit. Level 2 was scored when the internal organs also attached to the peritoneal wall, but it was still possible to free them. Level 3 was the extreme form of organ adhesion where the peritoneal lining tore apart and remained attached to the organs after the body cavity had to be forcibly opened. Note that peritoneal fibrosis levels 1, 2, and 3 used here correspond with levels 2, 3, and 4, respectively, used by Hund et al. (2020). For illustration of the 0-4 scale (Hund et al. 2020) see video at https://youtu.be/yKvcRVCSpWI. We also recorded total and dissected weight of the dead fish (0.001 g, Sartorius Element ELT202), fish sex (where it could be determined), and the presence of any internal parasites.

We excluded two species before data analysis. The clown feather-back (*Chitala ornata*) were already extremely fibrosed (level 3) when they arrived to our facility due to unknown reason and the kissing gourami (*Helostoma temminckii*) developed white spot disease during the experiment after they were injected. This made the total number of species tested 17 (Table 1).

### Data analysis

We formally tested the interaction between the effect of treatment (PBS, TH, Alum) and experimental species identity on individual fibrosis score using generalized least squares (GLS) model (function gls, library “nlme” v.3.1-148, Pinheiro et al. 2018). The response variable was fibrosis level (ordered integers 0-3) scored from the individual’s left flank. We set treatment, species and their interaction as fixed model effects. We attempted to originally analyze the ordinal response variable using Cumulative link models (CLM, function clm, library “ordinal” v.2019.12-10, Christensen 2019), but CLM with treatment–species interaction failed to converge due to model singularity (e.g., too many groups with zero variance because all individuals had identical fibrosis scores).

To assess phylogenetic signal in the experimental data, we created a species tree based on the recent comprehensive phylogeny of ray-finned fishes by Hughes et al. (2018) with estimated divergence times. We then obtained phylogenetic signal (i.e. the tendency of related species to show similar response) and ancestral character state for two species “traits”: the *maximum level* of peritoneal fibrosis in a species and *species relative response* in the positive control (Alum) compared to negative control (PBS). The *species relative response* accounts for some species having basic fibrosis level 1 while 0 was observed in the majority (12/17 species; Fig. 1). We measured phylogenetic signal with Pagel’s λ (function phylosig, with method specification to “lambda”, library “phytools” v.0.7-47 (Revell 2012)). We then estimated ancestral character state with maximum likelihood approach using function anc.ML (specifying Brownian motion mode of evolution, library “phytools”). All analyses were performed in R software v. 4.0.1 (R Core Developmental Team 2019).

**Figure 1.**
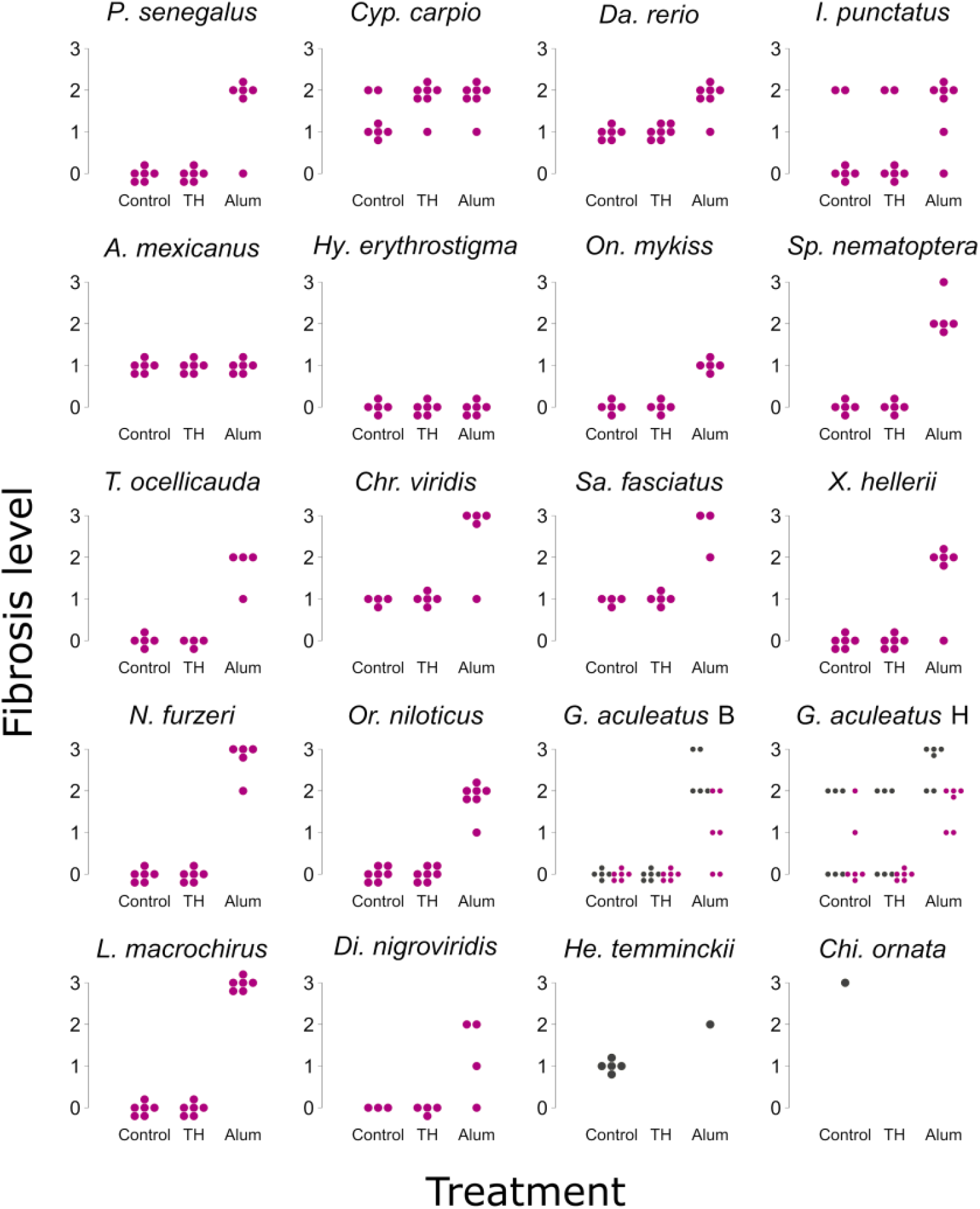
Peritoneal fibrosis in the experimental fish species. Individual points in the species plots show recorded level of peritoneal fibrosis scored from their left flank (the side of the injection). The fibrosis level was scored on an ordinal scale 0-3 and the jitter was used to show all data points. Note that the three-spine stickleback (both populations of *G. aculeatus*) that stayed in the experiment for 10 days are shown in grey as they did not enter data analysis. For completeness, we also present data for two unused species in grey - kissing gourami (*H. temminckii*) and clown feather-back (*Ch. ornata*). Full version of the abbreviated species names can be found in Table 1. The *G. aculeatus* B is for population from Loch a’ Bharpa and *G. aculeatus* H for Loch Hosta.

## RESULTS

### Literature review: Peritoneal fibrosis in fish is known but not well documented

Our first aim was to gather an overview of publication record encompassing peritoneal fibrosis in fish. In the articles we collected (for detailed methods see *Appendix*), general fibrotic response was documented from a wide array of ray-finned fish (Actinopterygii). We found 375 out of the 1335 articles (i.e. 28%) to be suitable for our study (i.e. articles mentioning fibrosis or encapsulation from Actinopterygii). The most-represented species came from the orders Cypriniformes (61 [including articles with multiple species]; 16% of relevant articles) and Salmoniformes (51; 14%) (Table 2). Authors typically identified signs of fibrosis during an autopsy, though some cases were observed from fish integument as well (e.g. capsules of skin parasites). Most of the suitable articles reported parasitism (197; 53%) or treatment (81; 22%) as the cause. Overall, fibrosis was present either on its own (175; 47%) or in combination with encapsulation of a foreign object (22; 6%).

**Table 2.**
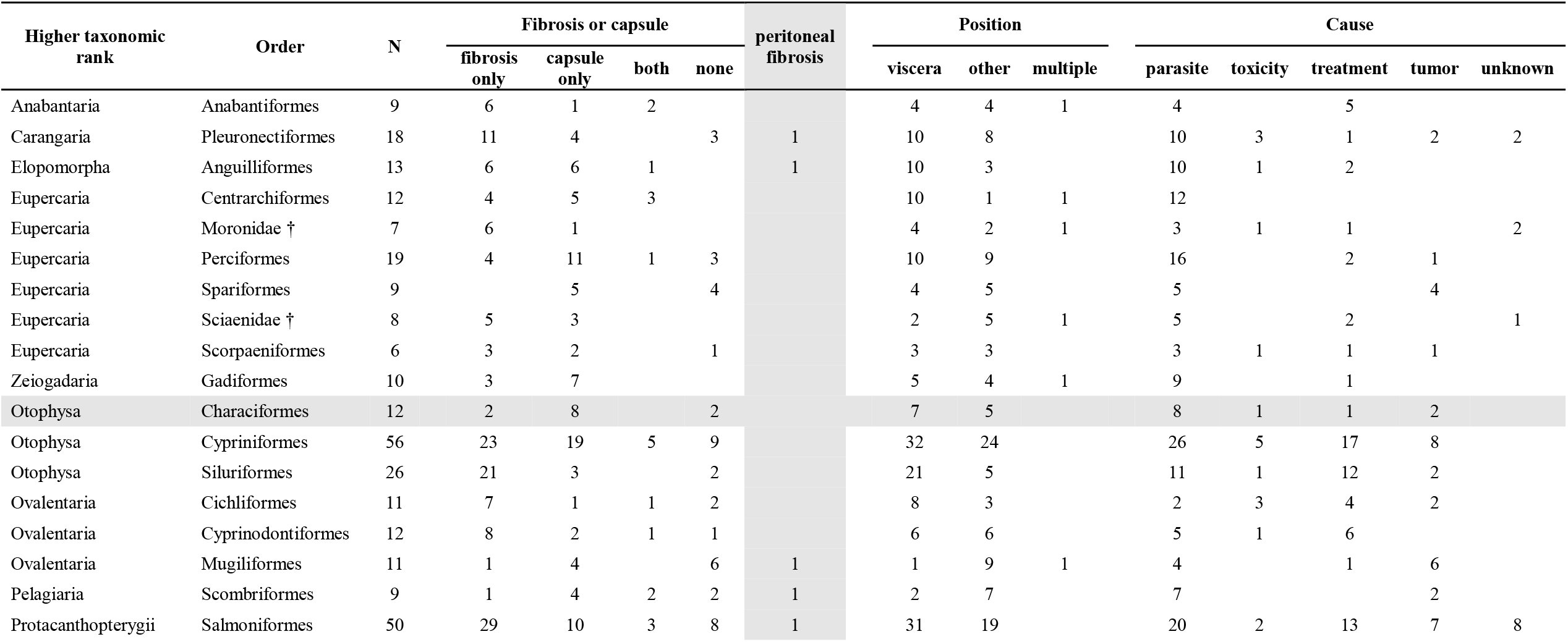
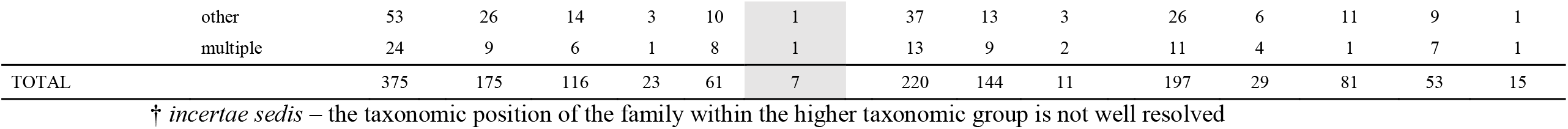
Overview of the literature search for presence of peritoneal fibrosis in fish. We show fish phylogenetic group, the presence of fibrosis and/or encapsulation, its location and cause. For brevity, we pooled less represented fish orders (with <6 articles) into “other” category and also grouped articles with species from more orders into “multiple”. Highlighted in gray are records of peritoneal fibrosis (column) and articles from Characiformes (row). Characiformes include the two tetra species that did not respond with peritoneal fibrosis to any of the treatments in our immune challenge experiment.

The extent of fibrosis was usually described only qualitatively, along with the identity of the affected organs or tissues. Taking articles related to fibrosis (reporting fibrosis alone or in combination with encapsulation, 198 articles), its incidence was mainly confined to visceral organs (152 cases, 77%). Fibrosis located around or in the internal organs was, however, mainly interstitial fibrosis often represented by tissue scarification after damage. We found only 7 articles that were dealing with peritoneal fibrosis specifically. These seven articles are diverse with regard to species taxonomic position and the cause of the fibrosis response (e.g., tapeworm infection, vaccination, radio-transmitter implantation) (Table 2). From this literature survey, we conclude that fibrosis is known in a number of fish species. Yet, the peritoneal fibrosis is very scarcely reported which limits our ability to draw broader conclusions about its evolutionary history or its function.

### Comparative immunological experiment: Common and strong immune response to Alum

We conducted a phylogenetically structured experimental study of ray-finned fish peritoneal fibrosis in response to immune challenge to reach more systematic understanding of its evolution. The level of fibrosis differed between experimental treatments and across species (GLS, treatment-species interaction, *F*_*34,251*_ = 7.93, *P* < 0.001) (Fig. 1). For most species, we observed no detectable or low fibrosis in control-injected (PBS) fish as well as in the tapeworm homogenate (TH) treatment (Fig. 1). However, there were two species, the common carp (*Cyprinus carpio*) and the channel catfish (*Ictalurus punctatus*), with variable individual response both in the PBS control and TH treatment (Fig. 1). Also note that in some species (like *D. rerio, S. fasciatus, C. viridis*), the default fibrosis level was 1 (organs attached together).

In contrast to the negative control and TH, the positive control (Alum injection) induced strong peritoneal fibrosis in most of the species (15 of 17). The response was typically high to extreme (fibrosis level 2 or 3, Fig. 1). The exceptions were two species of tetras, the Mexican tetra (*Astyanax mexicanus*) and the bleeding-heart tetra (*Hyphessobrycon erythrostigma*), which did not respond to any of the treatments (Fig. 1).

Phylogenetic signal for species maximum fibrosis and species relative response to the positive control (average Alum vs. PBS difference) both appeared to be strong, significantly different from random evolution (Pagel’s λ > 0.987 and *P* < 0.005, for both traits). Ancestral state at the base of the ray-finned fishes, i.e. at the divergence between the Senegal bichir (*Polypterus senegalus*) and the other species from our experiment, was estimated 1.862 for the species maximum fibrosis (Fig. 2) and 1.325 for species’ relative fibrosis response to the Alum treatment (average difference between Alum vs. PBS).

**Figure 2.**
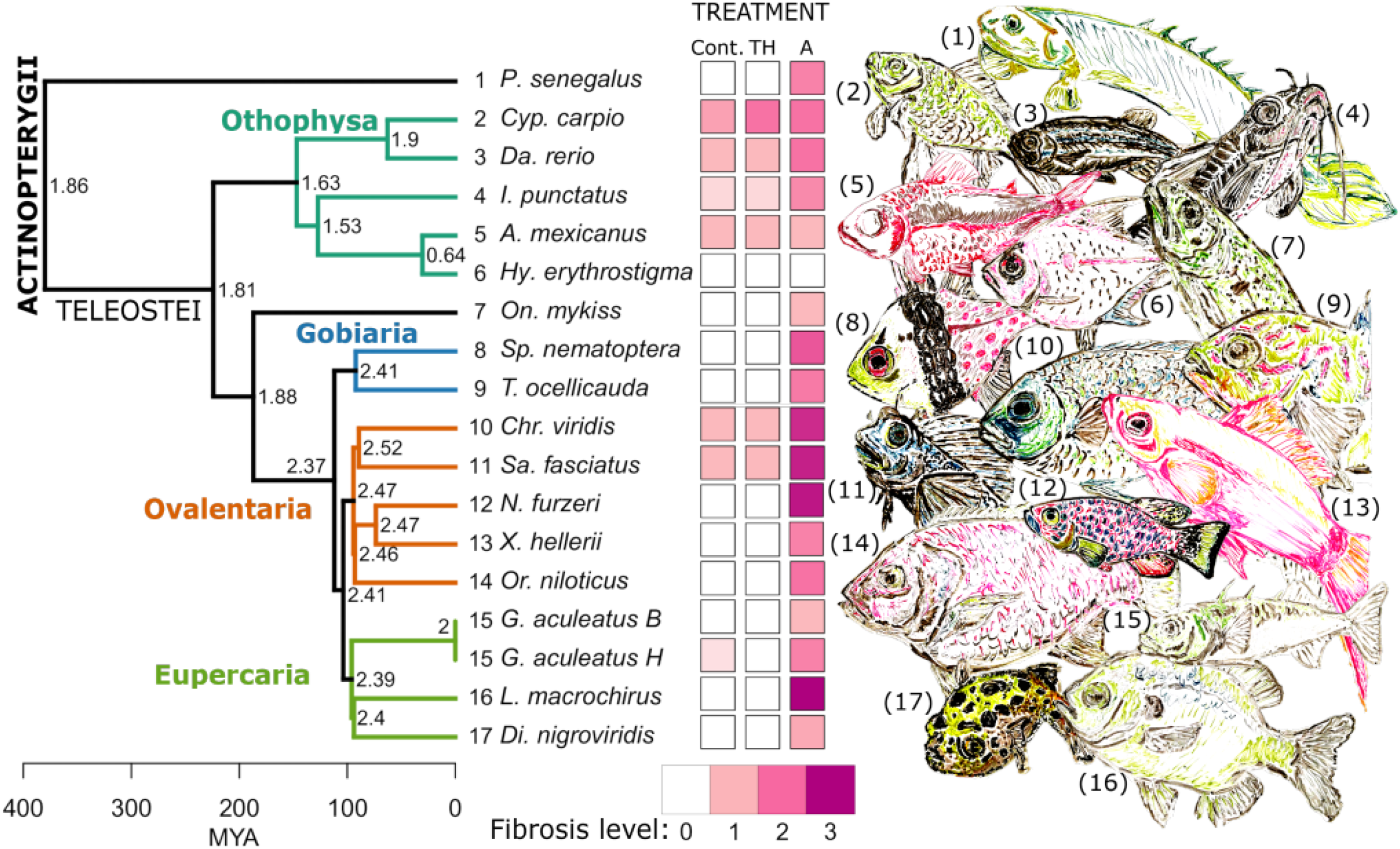
Ancestral state reconstruction of species maximum peritoneal fibrosis. The values at the branching nodes give estimates of the ancestral level of maximum peritoneal fibrosis level. Tree structure and branch lengths are based on recent reconstruction of phylogeny of ray-finned fishes (Actinopterygii) (Hughes et al. 2018). The squares of different intensity of purple colour show average level of peritoneal fibrosis per treatment in each tested species. Treatment abbreviations are Cont.: control (phosphate-buffered saline solution), TH: tapeworm antigen homogenate, A: Alum vaccine adjuvant. Fish species drawings by M. F. Maciejewski (not to scale). Full version of the abbreviated species names can be found in Table 1. The *G. aculeatus* B is for population from Loch a’ Bharpa and *G. aculeatus* H for Loch Hosta.

## DISCUSSION

This study represents one of the first comparative experimental assays of the macroevolution of an immune response. We focused on evaluating the prevalence of peritoneal fibrosis response across fishes, because this response has been shown to contribute to parasite growth suppression and elimination in some recently-evolved post-glacial lake populations of threespine stickleback. We show that published literature contains little data on peritoneal fibrosis in ray-finned fishes. To fill this gap, we experimentally tested the prevalence of peritoneal fibrosis in a wide array of species across the phylogeny of Actinopterygii. Our immune challenge resulted in a variable level of fibrosis between treatments and among species. Response to homogenate from the stickleback-specialized tapeworm was weak at best. This finding confirms that the use of fibrosis pathways in response to *S. solidus* is a recently-evolved trait. The positive control treatment (Alum), on the other hand, provoked strong peritoneal fibrosis in most of the species tested except one specific lineage – two species of tetras (from family Characidae, Characiformes). The results therefore suggest that, despite being rarely observed or reported, the capacity for peritoneal fibrosis is a phylogenetically widespread aspect of fish immunity. Ancestral character state reconstruction indicates that the peritoneal fibrotic response is phylogenetically conserved at least to the origin of ray-finned fishes (Fig. 2), estimated around 380 million years ago (Hughes et al. 2018).

### Lack of peritoneal fibrosis in publications across ray-finned fish

The literature search indicated that fish initiate fibrosis most frequently in response to tissue injury and/or parasitism. Thus, we can identify two main roles of fibrosis – maintenance of homeostasis in damaged tissue, and formation of physical barrier around an invader (encapsulation). Indeed, Gause et al. (2013) proposed an evolutionary hypothesis for the origin of parasite encapsulation from the ancestral repair response to tissue mechanical damage. The article collection also contained several cases (almost 1/3 of the articles mentioning fibrosis or encapsulation), where parasite encapsulation happened without more wide-spread fibrosis.

Our study was motivated by inter-population variation in threespine sticklebacks’ ability to encapsulate parasitic tapeworm *S. solidus* (Weber et al. 2017b). The inter-population variation probably stems from an evolutionary trade-off between the benefit of resistance (early encapsulation of the worm) and the risk of organ adhesion, excessive fibroblast proliferation, and ultimately partial sterility in the stickleback (Weber et al., in preparation; De Lisle and Bolnick 2020). Previous records on peritoneal fibrosis in threespine stickleback are mostly lacking (but see Hoffman 1975, p.175), although the phenomenon is found in numerous populations across the species’ circumpolar range. This is in line with the literature search, where we found only very few studies reporting peritoneal fibrosis in fish. In these few articles, peritoneal fibrosis was associated with serious intrusion of body integrity, e.g., radio-transmitter implantation (Mangan 1998), vaccination with bacteria (Colquhoun et al. 1998), or, similarly to the stickleback, tapeworm infection (Abdelsalam et al. 2016). Apparently, the stress has to be intensive and/or chronic to trigger peritoneal fibrosis. The question thus remained whether peritoneal fibrosis is that rare and highly specific response across fish species. We used a phylogenetically informed immune challenge experiment with selected representatives across the fish tree of life to answer this question.

### Peritoneal fibrosis in response to vaccination adjuvant was prevalent in most fish species

We successfully triggered peritoneal fibrosis in most of the fish species tested with the positive control (Alum). Alum is a commonly used vaccination adjuvant that causes influx of multiple types of immune cells into the injected region and alerts individual’s immune system (Kool et al. 2012). Yet, the particular mechanism of how Alum promotes vaccination is still unknown. In the case of peritoneal fibrosis, the Alum crystals may in fact act as an irritating agent that stimulates the type 2 immune response leading to containment of the body non-self (Gause et al. 2013). The widespread response to the Alum injection demonstrates that the capacity for peritoneal fibrosis is distributed across ray-finned fish phylogeny. Absence of fish peritoneal fibrosis in the published literature may thus stem from high specificity (peritoneal cavity invasion), low severity/chronicity of the common stressors, or the phenomenon may simply have been overlooked as was until recently in stickleback.

### The experimental exceptions

#### Individual variation

The homogenate prepared from *S. solidus* tapeworm caused peritoneal fibrosis only in two tested species and one population of the threespine stickleback. In common carp (*Cyprinus carpio*) and channel catfish (*Ictalurus punctatus*), the level of peritoneal fibrosis varied among individuals both in the TH and negative control (PBS) treatments. We recorded similar pattern also in threespine stickleback from Loch Hosta population after 10 days post-injection. The response was comparable between TH and the negative control (PBS). Based on the individual variation and the pre-treatment dissections, it seems like different individuals of the carp, catfish and Loch Hosta stickleback might be more or less sensitive to the injection itself. Individual variation in these three groups contrasts with the largely uniform response exhibited in each treatment by the remaining species.

#### General indifference to the TH treatment

*Schistocephalus solidus* is a parasite specialist of the threespine stickleback. The cue to trigger peritoneal fibrosis in threespine stickleback in response to the tapeworm is probably specific, as shown by some populations that fail to respond to certain genotypes of *S. solidus* (Weber et al. 2017a). Parasite community in North Uist stickleback is relatively rich and includes *S. solidus* (Rahn et al. 2016). The Scottish stickleback were also observed to show peritoneal fibrosis in the wild, with a’ Bharpa population having strong response and Hosta population absent (A. MacColl, pers. comm.). The tapeworm protein effective dose used here (9 mg/g of fish live weight) triggered strong fibrosis in a naturally fibrotic lake population from Vancouver Island (Hund et al. 2020), but it is possible that the Scottish stickleback do not recognize tapeworms from western Canada. The specific cue that triggers peritoneal fibrosis in response to the tapeworm infection is unknown, though presumably protein-based, and the work on its identification currently ongoing. It may still be as well possible that the expression of peritoneal fibrosis might be suppressed by stickleback in some non-fibrotic populations (Lohman et al. 2017; Fuess et al. 2020).

#### Absence of peritoneal fibrosis response

Our experimental data show that peritoneal fibrosis is widespread across fish phylogeny, except for two related species – the Mexican tetra (*Astyanax mexicanus*) and the bleeding-heart tetra (*Hyphessobrycon erythrostigma*), which did not respond to any of the treatments. The adaptation of some populations of Mexican tetra to the freshwater cave systems and lower parasitic burden could partially explain the observed pattern (Peuß et al. 2020). Taking into account the risks associated with the peritoneal fibrosis described in the threespine stickleback, maintenance of such response could be too costly for the Mexican tetra. Interestingly, the cave Mexican tetras exhibit marked reduction of visceral adipose tissue immunopathology compared to the surface populations (Peuß et al. 2020). However, we provide an evidence that the lack of the peritoneal fibrosis response might be more prevalent among Characiformes: both of the tetra species in our study didn’t respond to Alum. The non-specific immunity of these tetra species might therefore differ from the other ray-finned fish, and is a tempting target for more research, though we cannot yet generalize to many populations of each species, or across the entire clade. The family Characidae containing the two tetras consists of over a thousand species widely distributed in fresh waters from Texas, USA to Argentina. It would be interesting to uncover the mechanism for and phylogenetic extent of the absence of peritoneal fibrosis in more detail. Interestingly, our literature search indicates that Characiformes are able to encapsulate parasites and also show signs of (interstitial) tissue fibrosis (Table 2).

## Conclusion

As proposed by Gause et al. (2013), fibrosis is probably an ancient trait evolved from wound healing; the formerly repairing mechanism now suits also coping with endoparasites. We showed that, despite being rarely reported in the published literature, the potential to develop peritoneal fibrosis is widespread across fish phylogeny and it can be triggered through a general treatment (Alum peritoneal injection) in almost all tested fish. The comparative immunology experiment, such as the one we performed, is particularly powerful and broad approach to infer historical origin, evolutionary rate of the immune traits, and to identify interesting atypical lineages (Weber and Agrawal 2012). By investigating those exceptions, we may consequently focus on documenting genetic mechanisms and adaptive value of different character states.

## Author contributions

DIB and MV designed the study, MV conducted experimental work, performed data collection and analysis, MV and DIB wrote the manuscript.

## Acknowledgements

We would like to thank to Katherine R. Lewkowicz, Meghan F. Maciejewski, Lauren E. Fuess, Amanda K. Hund, Mariah L. Kenney, Foen Peng and Stephen P. De Lisle (members of the Bolnick Lab, University of Connecticut) for their help throughout the fish experiment and discussions on peritoneal fibrosis. Comments by Amanda K. Hund, Martin Reichard, Jakub Žák, Radim Blažek, Markéta Ondračková and Matej Polačik (Institute of Vertebrate Biology, CAS) helped to improve the manuscript. We are thankful to Quinebaug Valley State Fish Hatchery (CT, USA) for providing rainbow trout. MV’s stay at the University of Connecticut was supported by Fulbright Commission fellowship for research scholars. The project was funded by NIH project (NIAID grant 1R01AI123659-01A1) held by DIB. The experimental work was approved by University of Connecticut, Protocol No. A18-0008.

## Data Accessibility Statement

Data will be made publicly available after acceptance along with the statistical code under DOI: 10.6084/m9.figshare.12619367.

## Conflict of interest

The authors have declared no conflict of interest.

## APPENDIX

### Literature review on peritoneal fibrosis in fish - Methods

To gather an overview of publication record encompassing peritoneal fibrosis in fish, we performed two searches using Web of Science database on September 30, 2019. The first search was oriented towards parasite-induced fibrosis in fish with terms: (parasit^*^) AND (fibro^*^) AND ((teleost) OR (fish)). The other literature search was focused on fibrosis in fish in general while we tried to avoid articles on human subjects: (fibrosis AND (fish OR teleost) NOT (human)).

We collected 1459 entries in total, of which 1337 articles were retained after double entries removal. We considered an article to be suitable for our study if it was on ray-finned fish (Actinopterygii) and contained information on any signs related to fibrosis, like organ adhesion, spontaneous proliferation of fibroblasts (fibroma, fibrosarcoma), healing fibroplasia (scarification), or encapsulation (of a parasite or an implant, for example), that could be inferred from the title or article abstract. Encapsulation typically meant that an extra-bodily particle was surrounded with a layer of fibroblasts and extracellular matter (e.g., collagen fibers). We then sorted the suitable articles, by going into their main text, according to three criteria - the explicit presence of fibrosis or capsule (“fibrosis only”, “capsule only”, “both”, “none”), its topology (“viscera”, “other”, or “multiple”) and assumed cause (“parasite”, “toxicity”, “treatment”, “tumor”, or “unknown”). For topology (location) classification, we took internal organs related to excretory system, digestive tract, or reproduction (kidney, liver, gas bladder, gut, gonads, including also peritoneum) as “viscera” and the remaining organs or tissues, such as gills, skin, muscle, brain, heart, etc. as “other”. When tissues of both types were affected, we labelled the article as “multiple”. We then grouped articles with respect to the given cause of the fibrosis-related marks or encapsulation, where “parasite” was for infection with a uni- or multi-cellular parasite, “toxicity” was when the study monitored known environmental pollution (e.g., heavy metals), “treatment” was for deliberate manipulation with fish body or their living conditions (e.g., adding estrogen into water to test the effect on male physiology), “tumor” was when fibrosis happened spontaneously (e.g., fibrosarcoma) and “unknown” pooled studies where the cause could not be identified. We extracted fish species names and sorted them into orders and higher taxonomic categories according to the recent phylogenetic resolution of the Actinopterygii tree of life by Hughes et al. (2018) and Rabosky et al. (2018). We then used this dataset to offer an insight into the published literature on fish fibrosis.

## Notes

### Competing Interest Statement

The authors have declared no competing interest.

### Summary of Updates

In the revised version, we accommodated suggestions by referees and overall the changes are modest. We decided to short the manuscript and edit the figures for better comprehension.

## REFERENCES

Abdelsalam, M., R. Abdel-Gaber, M. A. Mahmoud, O. A. Mahdy, N. I. M. Khafaga, and M. Warda. 2016. Morphological, molecular and pathological appraisal of Callitetrarhynchus gracilis plerocerci (Lacistorhynchidae) infecting Atlantic little tunny (Euthynnus alletteratus) in Southeastern Mediterranean. J. Adv. Res. 7:317–326.

Allen, J. E., and R. M. Maizels. 2011. Diversity and dialogue in immunity to helminths. Nat. Rev. Immunol. 11:375–388.

Arme, C. and R.W. Owen. 1967. Infections of the three-spined stickleback, Gasterosteus aculeatus L., with the plerocercoid larvae of Schistocephalus solidus (Müller, 1776), with special reference to pathological effects. Parasitology, 57:301–314.

Barber, I., and J. P. Scharsack. 2010. The three-spined stickleback-Schistocephalus solidus system: An experimental model for investigating host-parasite interactions in fish. Parasitology 137:411–424.

Colquhoun, D. J., E. Skjerve, and T. T. Poppe. 1998. Pseudomonas fluorescens, infectious pancreatic necrosis virus and environmental stress as potential factors in the development of vaccine related adhesions in Atlantic Salmon, Salmo salar L. J. Fish Dis. 21:355–364.

Christensen, R. H. B. 2019. ordinal - Regression Models for Ordinal Data. R package version 2019.12-10. https://CRAN.R-project.org/package=ordinal.

De Lisle, S. P., and D. I. Bolnick. 2020. Reproductive fitness costs of an immune response in the wild. bioRxiv 1–40. DOI: 10.1101/2020.07.10.197210.

Flajnik, M. F., and L. Du Pasquier. 2004. Evolution of innate and adaptive immunity: Can we draw a line? Trends Immunol. 25:640–644.

Fuess, L., J. N. Weber, S. Den Haan, N. C. Steinel, K. C. Shim, and D. I. Bolnick. 2020. A test of the Baldwin Effect: Differences in both constitutive expression and inducible responses to parasites underlie variation in host response to a parasite. bioRxiv: 1–30. DOI: 10.1101/2020.07.29.216531.

Gause, W. C., T. A. Wynn, and J. E. Allen. 2013. Type 2 immunity and wound healing: Evolutionary refinement of adaptive immunity by helminths. Nat. Rev. Immunol. 13:607– 614.

Herpin, A., C. Lelong, and P. Favrel. 2004. Transforming growth factor-β-related proteins: An ancestral and widespread superfamily of cytokines in metazoans. Dev. Comp. Immunol. 28:461–485.

Hoffman, G. L. 1975. Lesions due to internal helminths of freshwater fishes, pp. 151–188. in Ribelin, W. E., and G. Migaki (eds.). The Pathology of Fishes. University of Wisconsin Press, Madison, WI, USA.

Hughes, L. C., G. Ortí, Y. Huang, Y. Sun, C. C. Baldwin, A. W. Thompson, D. Arcila, R. Betancur-Rodriguez, C. Li, L. Becker, N. Bellora, X. Zhao, X. Li, M. Wang, C. Fang, B. Xie, Z. Zhoui, H. Huang, S. Chen, B. Venkatesh, and Q. Shi. 2018. Comprehensive phylogeny of ray-finned fishes (Actinopterygii) based on transcriptomic and genomic data. Proc. Natl. Acad. Sci. 115:6249–6254.

Hund, A. K., L. E. Fuess, M. L. Kenney, M. F. Maciejewski, J. M. Marini, K. C. Shim, and D. I. Bolnick. 2020. Rapid evolution of parasite resistance via improved recognition and accelerated immune activation and deactivation. bioRxiv 1–35. DOI: 10.1101/2020.07.03.186569.

Kool, M., K. Fierens, and B. N. Lambrecht. 2012. Alum adjuvant: Some of the tricks of the oldest adjuvant. J. Med. Microbiol. 61:927–934.

Lazzaro, B., and A. Clark. 2012. Rapid evolution of innate immune response genes. Rapidly evolving genes and genetic systems, 203-222. in Singh, R. S., J. Xu, and R. J. Kulathinal (eds.). Rapidly evolving genes and genetic systems. Oxford University Press, Oxford, UK.

Litman, G. W., J. P. Cannon, and L. J. Dishaw. 2005. Reconstructing immune phylogeny: New perspectives. Nat. Rev. Immunol. 5:866–879.

Litman, G. W., and M. D. Cooper. 2007. Why study the evolution of immunity? Nat. Immunol. 8:547–548.

Lohman, B. K., N. C. Steinel, J. N. Weber, and D. I. Bolnick. 2017. Gene expression contributes to the recent evolution of host resistance in a model host parasite system. Front. Immunol. 8.

Malmstrøm, M., S. Jentoft, T. F. Gregers, and K. S. Jakobsen. 2013. Unraveling the evolution of the Atlantic cod’s (Gadus morhua L.) alternative immune strategy. PLoS One 8:1–9.

Malmstrøm, M., M. Matschiner, O.K. Tørresen, B. Star, L. G. Snipen, T. F. Hansen, and S. Jentoft. 2016. Evolution of the immune system influences speciation rates in teleost fishes. Nat. Genet. 48:1204–1210.

Mangan, B. P. 1998. Long-term retention of a radio transmitter by a muskellunge. J. Freshw. Ecol. 13:485–487.

Peuß, R., A. C. Box, S. Chen, Y. Wang, D. Tsuchiya, J. L. Persons, A. Kenzior, E. Maldonado, J. Krishnan, J. P. Scharsack, B. D. Slaughter, and N. Rohner. 2020. Adaptation to low parasite abundance affects immune investment and immunopathological responses of cavefish. Nat. Ecol. Evol. 4:1416–1430.

Pinheiro, J., D. Bates, S. DebRoy, D. Sarkar, and R Core Team. 2018. nlme: Linear and nonlinear mixed effects models. R package version 3.1-137. https://CRAN.R-project.org/package=nlme

Polačik, M., R. Blažek, and M. Reichard. 2016. Laboratory breeding of the short-lived annual killifish Nothobranchius furzeri. Nat. Protoc. 11:1396–1413.

R Core Team. 2020. R: A language and environment for statistical computing. R Foundation for Statistical Computing, Vienna, Austria. https://www.R-project.org/

Rabosky, D. L., J. Chang, P. O. Title, P. F. Cowman, L. Sallan, M. Friedman, K. Kaschner, C. Garilao, T. J. Near, M. Coll, and M. E. Alfaro. 2018. An inverse latitudinal gradient in speciation rate for marine fishes. Nature 559:392–395.

Radwan, J., W. Babik, J. Kaufman, T. L. Lenz, and J. Winternitz. 2020. Advances in the evolutionary understanding of MHC polymorphism. Trends Genet. 36:298–311.

Rahn, A. K., E. Eßer, S. Reher, F. Ihlow, A. D. C. MacColl, and T. C. M. Bakker. 2016. Distribution of common stickleback parasites on North Uist, Scotland, in relation to ecology and host traits. Zoology 119:395–402.

Revell, L. J. 2012. phytools: An R package for phylogenetic comparative biology (and other things). Methods Ecol. Evol. 3:217–223.

Rijkers, G. T., E. M. Frederix-Wolters, and W. B. van Muiswinkel. 1980. The immune system of cyprinid fish. Kinetics and temperature dependence of antibody-producing cells in carp (Cyprinus carpio). Immunology, 41:91–97.

Ritter, M., M. Kalbe, and T. Henrich. 2017. Virulence in the three-spined stickleback specific parasite Schistocephalus solidus is inherited additively. Exp. Parasitol. 180:133–140.

Simon, V., R. Elleboode, K. Mahé, L. Legendre, P. Ornelas-Garcia, L. Espinasa, and S. Rétaux. 2017. Comparing growth in surface and cave morphs of the species Astyanax mexicanus: Insights from scales. Evodevo 8:23.

Slodkowicz, G., and N. Goldman. 2020. Integrated structural and evolutionary analysis reveals common mechanisms underlying adaptive evolution in mammals. Proc. Natl. Acad. Sci. 117:5977–5986.

Solbakken, M. H., K. L. Voje, K. S. Jakobsen, and S. Jentoft. 2017. Linking species habitat and past palaeoclimatic events to evolution of the teleost innate immune system. Proc. R. Soc. B Biol. Sci. 284.

Thannickal, V. J., Y. Zhou, A. Gaggar, and S. R. Duncan. 2014. Fibrosis: ultimate and proximate causes. J. Clin. Invest. 124:4673–4677.

Velová, H., M. W. Gutowska-Ding, D. W. Burt, and M. Vinkler. 2018. Toll-like receptor evolution in birds: Gene duplication, pseudogenization, and diversifying selection. Mol. Biol. Evol. 35:2170–2184.

Weber, J. N., M. Kalbe, K. C. Shim, N. I. Erin, N. C. Steinel, L. Ma, and D. I. Bolnick. 2017a. Resist globally, infect locally: A transcontinental test of adaptation by stickleback and their tapeworm parasite. Am. Nat. 189:43–57.

Weber, J. N., N. C. Steinel, K. C. Shim, and D. I. Bolnick. 2017b. Recent evolution of extreme cestode growth suppression by a vertebrate host. Proc. Natl. Acad. Sci. 114:6575–6580.

Weber, M. G., and A. A. Agrawal. 2012. Phylogeny, ecology, and the coupling of comparative and experimental approaches. Trends Ecol. Evol. 27:394–403.

Weber, J., N. C. Steinel, K. C. Shim, D. Rennison, F. Peng, S. P. De Lisle, L. E. Fuess, and D. I. Bolnick. An evolutionary Pyrrhic victory via a costly but effective immune response to control cestode infection. In Prep.

